# SC10X/U: A High-density Electrode System for Non-Invasive Recording of Neural Activity of the Cervical Spinal Cord

**DOI:** 10.1101/2024.10.23.619533

**Authors:** Prabhav Mehra, Marjorie Metzger, Saroj Bista, Serena Plaitano, Eileen R. Giglia, Leah Nash, Éanna Mac Domhnaill, Matthew Mitchell, Peter Bede, Madeleine Lowery, Muthuraman Muthuraman, Orla Hardiman, Bahman Nasseroleslami

## Abstract

**Objective:** To design and develop a high-density (HD) electrode system that describes the position of surface electrodes for recording electrophysiological signals from the human cervical spinal cord. The system is intended to standardize experimental recordings and facilitate the subsequent analysis of evoked and spontaneous spinal cord neural activity, using high-density electrospinography (HD-ESG).

**Method:** The proposed system (SC10-X/U) describes the locations of up to 76 channels with a unique nomenclature, where the division of the spinal cord (SC) electrode space was inspired by the EEG 10-10 system. As proof of concept, spinal evoked potentials in response to median nerve stimulation at the wrist were recorded from 10 participants and characterized based on a 64-channel derivation from the SC10-X/U system.

**Results:** Following the design criteria, the SC10-X/U defines 76 electrode positions and its configuration. HD-ESG system was utilized to successfully record evoked spinal responses and significant N13 and P9 potential were observed in response to the stimulation. The spinal N13 potential had a latency of 13.2 +-1.1ms (mean +-SD) after stimulation. A topographic map of the N13 electro-spinal activity using the 64-channel recording system revealed an epicentre at C5-C7 dorsal-vertebral locations (ML4 - ML6 electrodes).

**Conclusion:** The proposed SC10-X/U system will facilitate standardized recording and analysis of high-density ESG signals from the human cervical spinal cord. The system defines electrode locations to promote standardization across different individuals, studies, and clinical and research centres. The HD-ESG evoked potentials recorded using the proposed system were comparable to those observed in previous non-HD studies. The presented topographic maps conform to known neurophysiological and neuroanatomical findings. This served to validate the design and development of the electrode system and patch for future studies.

## 1. Introduction

The spinal cord is an integral part of the sensorimotor nervous system and plays a crucial role in the neural communication between the cortical/sub-cortical regions and periphery, which is underpinned by complex neural circuits within the spinal cord (Pierrot-Deseilligny and Burke, 2012). These neural circuits are responsible for transmitting and processing sensorimotor signals and for establishing functional connections (Hochman, 2007). Although the SC plays an essential role in communication with sensorimotor networks, neurophysiological and imaging studies have largely treated it as a wire that relays information between the brain and the periphery, ignoring the neural complexity of the spinal cord. Similarly, neuroelectric imaging studies on sensorimotor communication have primarily focused on brain-muscle communication using Electroencephalography (EEG) – Electromyography (EMG) based analysis (Bao et al., 2021; Bista et al., 2023; Boonstra, 2013; Coffey et al., 2021; Conway et al., 1995); hence limiting our understanding of the neuro-electrophysiology of the role of spinal cord.

Non-invasive recording of spinal neural activity is necessary to assess and analyse the function and dysfunction of spinal cord in neurodegenerative diseases (e.g. Amyotrophic Lateral Sclerosis, ALS) and other neurological conditions (such as stroke), as well as across different physiological conditions in the healthy population at large. The use of spinal functional magnetic resonance imaging (fMRI) in spinal cord assessment is limited due to the structure and location of the spinal cord (Bede et al., 2012; Bede and Hardiman, 2014; Cohen-Adad, 2017). Non-invasive neuroelectric and neuro-electromagnetic imaging techniques provide high temporal resolution that is necessary for assessing spinal cord function, communication, and fast neural oscillations (Bankim Subhash Chander et al., 2022; Pierrot-Deseilligny and Burke, 2012).

In earlier electrophysiological studies, the function of the spinal cord was assessed by recording evoked potentials at lower cervical vertebral levels in response to electrical stimulation (Cioni and Meglio, 1986; Desmedt and Huy, 1984; Fujimoto et al., 2001; Prestor et al., 1997; Shimoji et al., 1971). More specifically, both invasive (Ertekin, 1976; Insola et al., 2008; Jeanmonod et al., 1991; Kaneko et al., 1998; Prestor and Golob, 1999) and non-invasive (Emerson et al., 1984; Restuccia and Mauguière, 1991; Schabrun et al., 2012; Zanette et al., 1995) recording methodologies were used to record somatosensory evoked potentials (SEPs) at spinal level in response to peripheral nerve stimulation to either understand spinal (dys)function or develop clinical biomarkers. Non-invasive short latency cervical spinal SEP’s at C6 vertebrae (Cv6) level were reported at average latencies of 9 ms (P9), 11 ms (N11), and 13 ms (N13) from the posterior region (Cruccu et al., 2008; Mauguière, 2003). The N13 potentials are proposed to be generated by post-synaptic signalling in the dorsal horn of the spinal cord (Desmedt and Cheron, 1981; Jeanmonod et al., 1989; Urasaki et al., 1990) and are considered to be the surface-level representation of the N1 potential observed in the P1-N1-P2 triphasic response via invasive epidural recording at the cervical spinal (Cioni and Meglio, 1986; Ertekin, 1976; Shimoji et al., 1971).

The earlier electrophysiological studies have primarily relied only on a limited number of channels to record the spinal activity from surface or from epidural sites. In the early 1980s, ring electrode placement system (Desmedt and Huy, 1984; Emerson et al., 1984; Restuccia and Mauguière, 1991) was proposed to record spatial spinal evoked activity at a pre-selected cervical vertebral level. The ring electrode placement standardized the system for recording both posterior and anterior spinal activity from a particular cervical vertebra. This system still remains an active standard to record the neuro-electric activity from a select lower cervical level that has been used in more recent studies (Bankim Subhash Chander et al., 2022).

Advancements in technology have increased our capability to record neural activity from a large number of channels as observed by the increase in spatial resolution of high-density (HD) EEG and EMG recordings. Similarly, multi-channel non-invasive recording of spinal activity have become more common to increase the spatial resolution. Recent studies of HD-Magnetospinography (MSG) (Akaza et al., 2020; Mardell et al., 2022; Sumiya et al., 2017) and “multi-channel electrospinography (ESG)” (Nierula et al., 2022) exploited multiple electrodes to record spinal activity, with the number of electrodes ranging from 16 to 44. Despite the rise of multi-channel spinal studies, there is no standardized electrode placement system for localizing the electrode space with respect to the spinal space. The absence of a recognized electrode placement standard has resulted in considerable variation in electrode locations across studies (Akaza et al., 2020; Mardell et al., 2022; Nierula et al., 2022).

In the late 1950s, the need for a standardized electrode location for EEG recordings was recognized, and the 10-20 electrode system was proposed by Jasper, HH. (H, 1958). This later advanced to the 10-10 (Chatrian et al., 1985) and 10-5 system (Oostenveld and Praamstra, 2001)as the resolution of EEG recording increased (Jurcak et al., 2007). We believe that high-density ESG will substantially benefit from a similar recognized placement system: a standardized electrode placement system that allows for compatible, comparable, and reproducible methods across research groups, participant groups, and clinicians. Moreover, this would promote standard practices that can be combined across large-scale neurophysiological and clinical studies on neurological conditions.

In this paper, we propose a high-density electrode placement system for non-invasive recording of neuroelectric signals from cervical and upper thoracic spinal cord. The proposed electrode placement system (SC10-X/U) can incorporate up to 76 channels aimed at standardizing electrode space with respect to the spinal space. This system was realised using a flexible patch for holding the electrodes. The utility of the SC10-X/U placement system is demonstrated by recording and analysing the evoked spinal potentials in response to median nerve stimulation from a 64-channel HD-ESG montage derived from the proposed SC10X/U system.

## 2. The SC10-X/U Electrode System

There is currently no recognized standard electrode placement system for recording electrical activity over the cervical and upper thoracic spinal cord. The proposed SC10-X/U is described in this section, following the statement of the design requirements and criteria. The SC10-X/U system can incorporate up to 76 channel electrodes for non-invasively recording spinal activity.

We used the basic principles of the EEG 10-10 system and implemented it in the spinal cord space to design a high-density ESG electrode placement system, based on the rationale that the underlying principles of the electrode placement system defined for the brain can also be applied to the spinal cord. The principles include defining a 2D representation of the region of interest (e.g., brain, spinal cord), finding reference anatomical landmarks, and diving the 2D space in proportional distances. In the EEG 10-10 system the 2D space is represented as a circle; anatomical references include nasion, inion, left and right pre-auricular points (LPA and RPA); and the space is divided into 10% proportional distances. These principles are extended to the spinal space, resulting in the electrode placement for spinal space. The electrode placement system is named SC10X/U system.

### 2.1. The Design Criteria and Principles of ESG electrode placement

#### 1. Two-dimensional representation of the space

The cervical-thoracic spinal space can be envisioned as two curved cylinders joined together. The cervical curvature (lordotic) and the thoracic curvature (kyphotic) can be characterized as concave and convex respectively, as shown in Figure *1*. The 2D representation of the curved surface of the cylinders is described by a rectangle with curved top and bottom.

**Figure 1:**
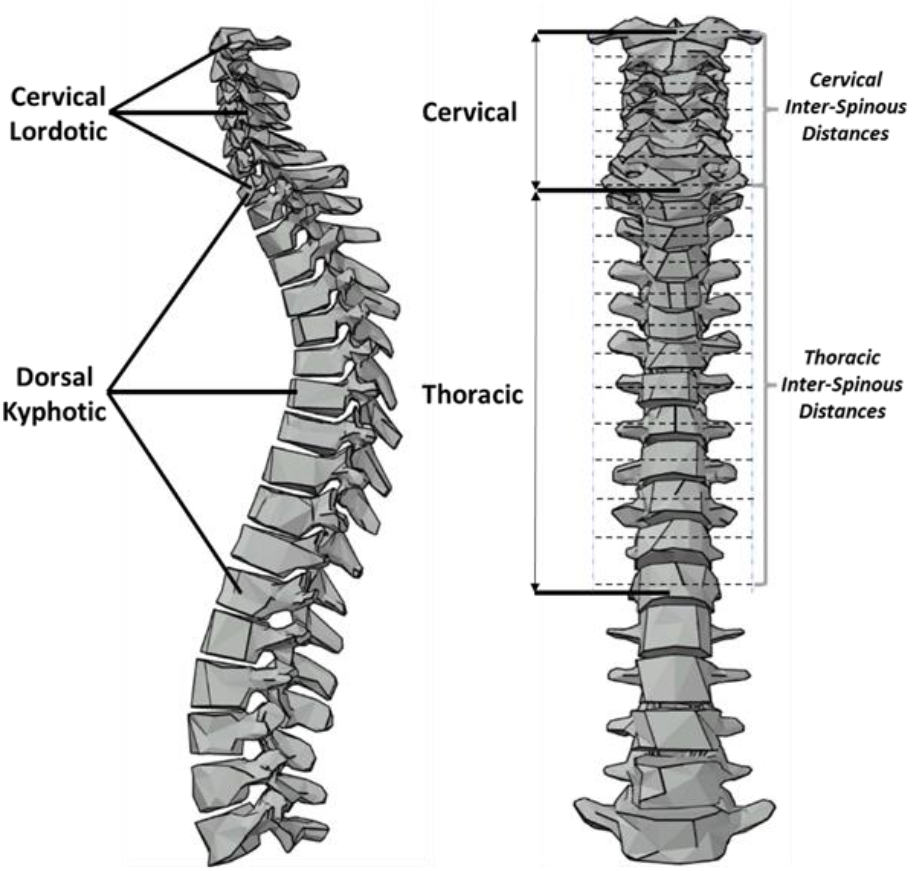
Spinal Cord CAD model of the cervical and thoracic regions. The left figure shows lordotic and kyphotic curvatures of the spinal cord. The right figure represents inter-vertebral spinous distances.

#### 2. Anatomical Landmarks

A total of 16 anatomical references for representing cervical and upper thoracic spinal space were selected. These include the occipital protrusion, C2 spinous process, cervical (C5 and C6) and thoracic (T1) spinous processes, left and right pre-auricular points, transverse spinous process of atlas (left and right), carotid tubercules (left and right), root of scapula (left and right), intersection of transverse plane through C5 spinous process and sagittal plane passing through inferior angle of scapula (left and right). These anatomical points encompass the cervical and upper thoracic spinal space as shown in Figure 2.

**Figure 2:**
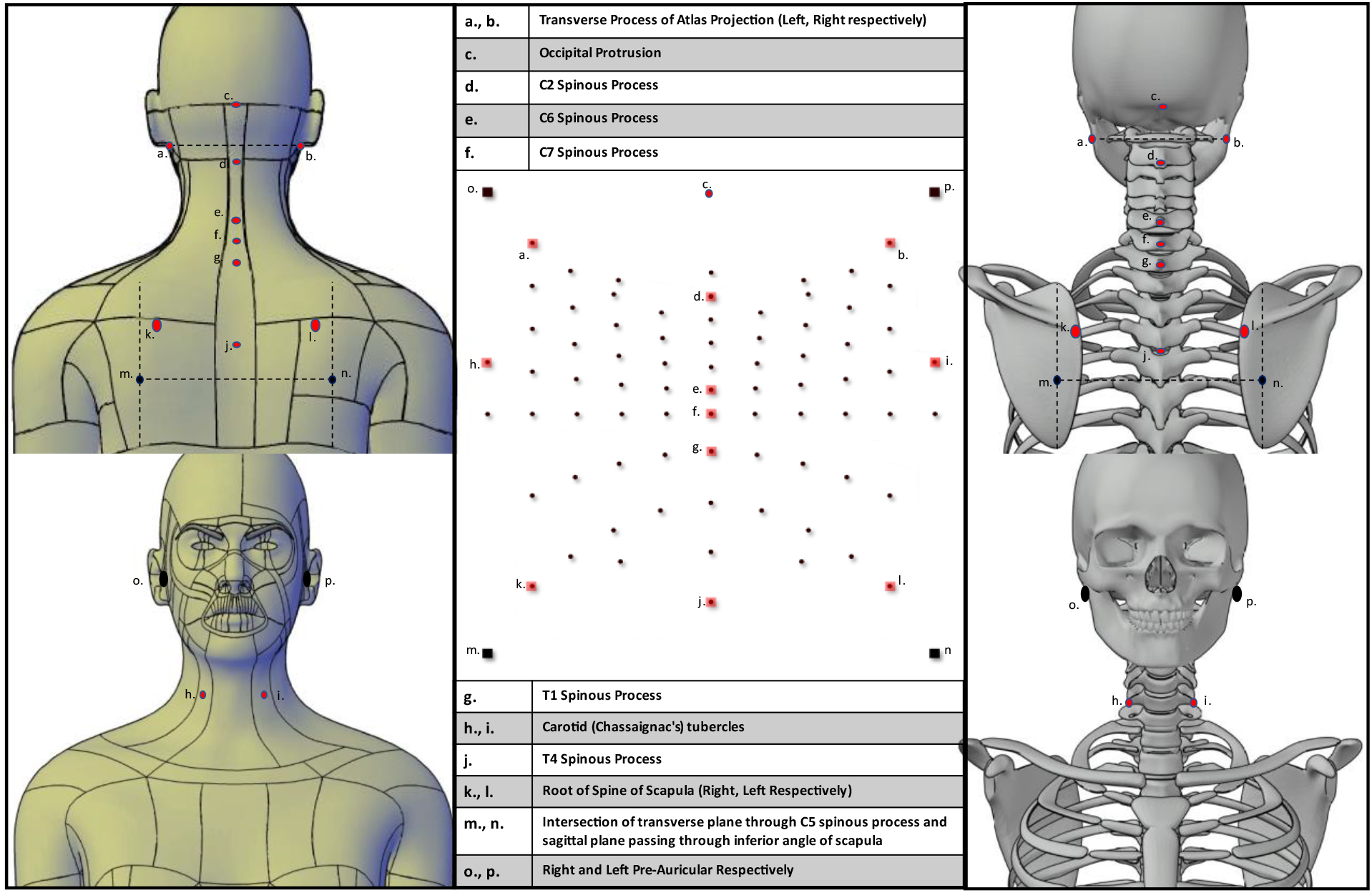
Anatomical References selected as surface landmarks are shown on the human model, skeleton model, and the SC10-X/U Electrode Placement System. The anatomical points encompassing spinal space are labelled from a. to p. and are shown over the surface for the human model (left), skeleton model (right), and electrode system (centre).

#### 3. Division of 2D space

The medial-transverse space is divided into 10 equal segments each accounting for 10% of the total lateral-medial-lateral distance, Figure 3. The medial-sagittal space along the length and direction of the spinal cord is divided in proportion to the inter-vertebrae spinous distances contained in the electrode space (Figure *1*) shows the inter-vertebral spinous distances). The medial sagittal space encompassing the cervical spine region is divided in proportion to the average cervical intervertebral spinous distance relative to the total medial sagittal distance. Similarly, the space that encompasses the thoracic region is divided in proportion to the average thoracic intervertebral spinous distance relative to the total medial sagittal distance. The division of medial sagittal and transverse space is highlighted in Figure 3.

**Figure 3:**
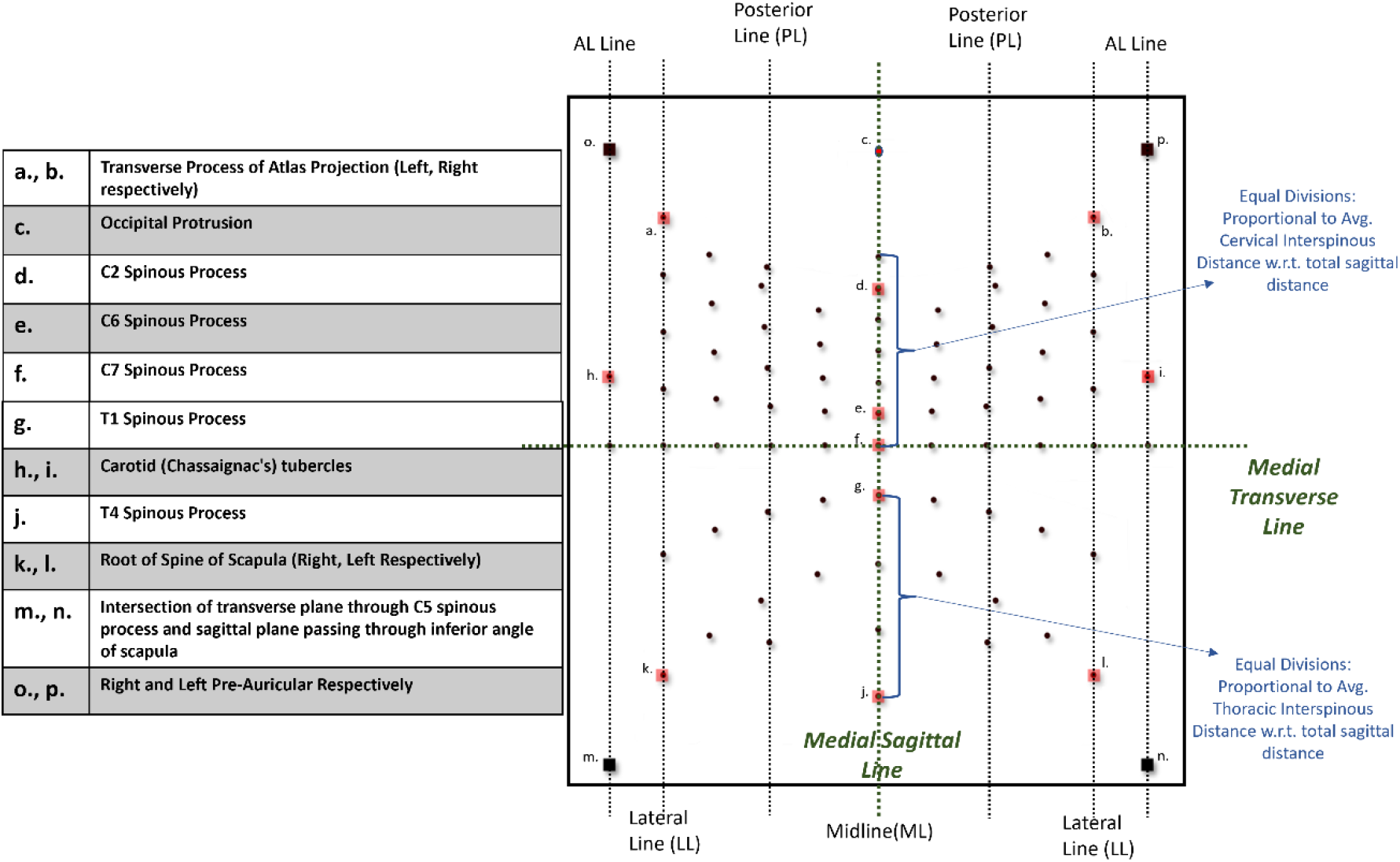
Electrode placement system (SC10-X/U) representing spinal space for high-density surface electrospinography (HD-ESG). The SC10-X/U system for the 76-electrode locations is shown along with anatomical reference points and spatial features, which are utilized for 2D space division.

### 2.2. Design and Characterisation of the SC10-X/U Electrode Placement System

The electrode placement system shown in Figure 3 is designed to cover the cervical and upper thoracic spinal cord space for surface neuro-electro-physiological recordings. The boundary of the space is based upon the anatomical reference points to standardize the electrode placement system. The upper boundary points are defined by the right pre-auricular and left pre-auricular points (LPA and RPA). The bottom boundary points of thoracic region are defined by the intersection of transverse plane at T5 spinous level and sagittal planes passing through the inferior angle of scapula (marked as m. and n., Figure 2). The lateral end points of the cervical region are defined by left and right C6 anterior tubercles (marked as h. and i., Figure 2). The boundary is thereby defined by connecting the marked boundary points as shown in Figure 4.a. The transverse plane passing through the C7 spinous process is divided in 10% proportional segments (ten equidistance points) between left and right lateral ends as shown in Figure 3 and Figure 4.

**Figure 4:**
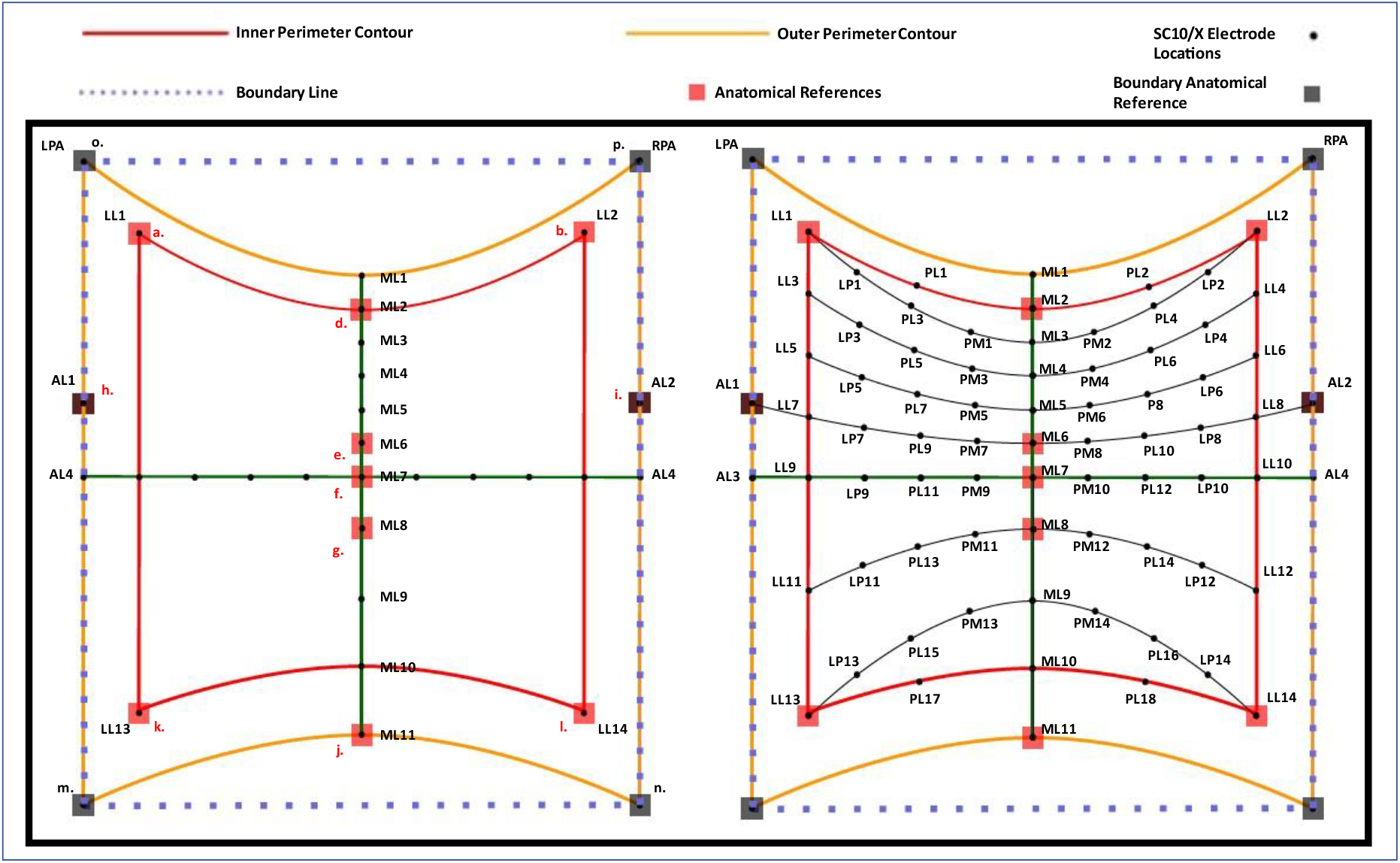
Design description of the SC10-X/U electrode placement system with electrode locations and labels (right). The left figure showcases the reference boundary, inner and outer perimeter contour. The right figure showcases all 76-electrode locations and labels. The labels are defined as, ML: Midline, PM: Posterior Midline, PL: Posterior Line, LP: Lateral Posterior, LL: Lateral Line, AL: Anterior Line

**Figure 5:**
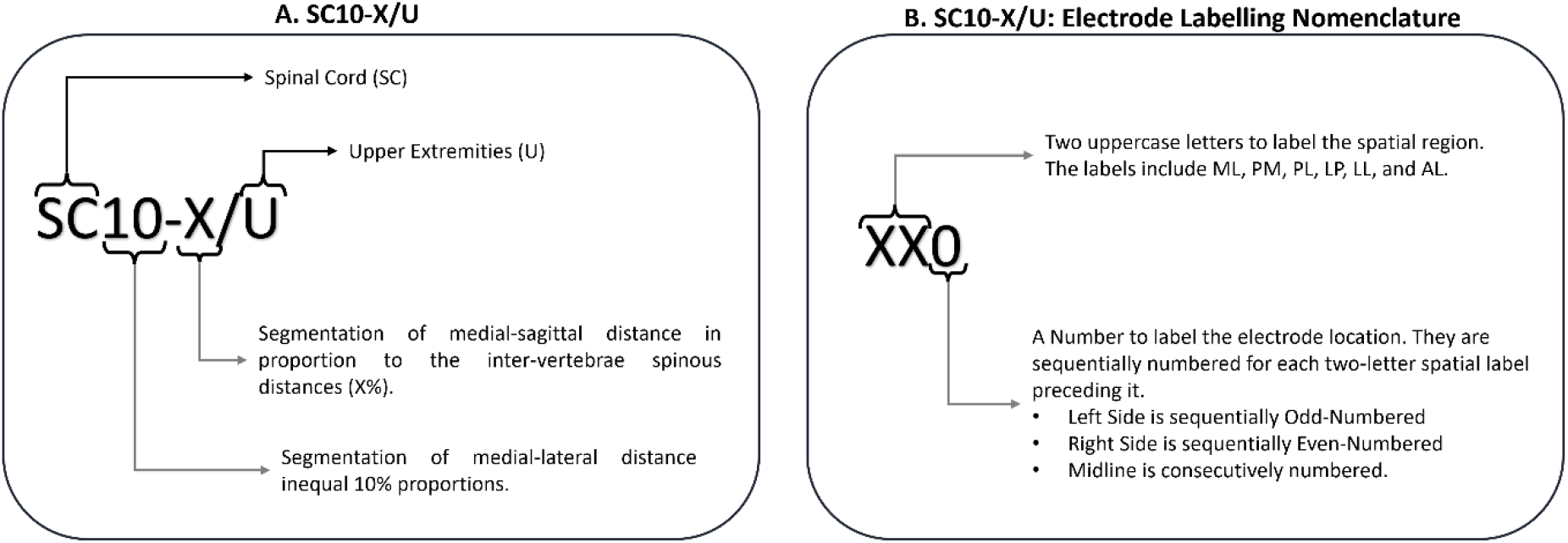
A. The SC10-X/U electrode placement system acronym. B. The SC10-X/U electrode location labelling nomenclature.

A line is subsequently drawn along the C2, C7, T1, and T4 spinous processes to define the midline (ML). The midline running along the spinal cord is divided into ten segments resulting in electrode positions labelled ML1 to ML11 (Figure 4.a). The inter-electrode distance between cervical midline electrodes (ML1-ML7) and upper thoracic midline electrodes (ML8-ML11) is in proportion to the average cervical inter-vertebral spinous distances and average thoracic inter-vertebral spinous distances relative to the total medial-sagittal distance (refer to figure 3 and figure 1). The midline between ML1 and ML7 spanning cervical vertebrae levels are divided in equal proportions corresponding to the average cervical inter-spinous distance. Similarly, ML8 and ML11 spanning upper thoracic vertebral levels are divided in equal proportions corresponding to the average thoracic inter-spinous distance (Figure 4.a). As a result, the defined electrode positions (ML1-ML11) along the midline roughly align with their respective vertebral levels.

Furthermore, the outer perimeter contour is defined by connecting LPA, RPA, ML1, ML11 position markers and AL lines (defined by connecting pre-auricular points, m., n., and C6 anterior tubercles on either side), encompassing the complete electrode space. The electrode positions one level below ML1, one level above ML11, and one level beside RPA and LPA on the lateral plane are marked as ML2, ML10, LL1, and LL2 respectively. This defines the inner-perimeter contour (Figure 4.a). The electrodes LL1 and LL2 reside at the level of the transverse process of the atlas (marked as a. and b., figure 2). The left and right lines of the inner-perimeter contour are defined as lateral lines (LL). The lateral line (LL) on the either side of contour is divided into six segments, four equal segments on cervical side and two equal segments on the thoracic side. The top and bottom segment of the left contour connecting LL1 to ML2 and LL13 to ML10 respectively are divided in two halves each. Similar divisions are made on the right top and bottom segment connecting LL2 to ML2 and LL14 to ML10. This division result in ten divisions on each half of the inner-perimeter contour. The division of inner-perimeter contour define the electrode position markers, labelled LL1, PL1, ML2, PL2, LL2, LL4, LL6, LL8, LL10, LL12, LL14, PL18, ML10, PL17, LL13, LL11, LL9, LL7, LL5, and LL3 as shown in Figure 4.b.

Finally, a transverse curve is defined by connecting LL1 to LL2 and passing through the midline at ML3 (LL1-ML3-LL2). The defined transverse curve is divided in 8 equal fractions, resulting in positions marked by LP1, PL3, PM1, PM2, PL4, and LP2 as shown in Figure 4.b. This is achieved by bisecting the curve twice on either side of the midline, dividing it into halves and quarters. Similar transverse curve are determined along the electrode markers LL3-ML4-LL4, LL5-ML5-LL6, LL7-ML6-LL8, LL9-ML7-LL10, LL11-ML8-LL12, and LL13-ML10-LL14 positions. These transverse curves are similarly divided in 1/8 fractions to define the remaining electrode position markers.

These steps define the resultant SC10-X/U electrode placement system (Figure 4b) determined to cover the spinal space encompassing cervical and upper thoracic spinal cord.

### 2.3. SC10-X/U Electrode Labels Nomenclature

The labels used in the SC10-X/U nomenclature for the electrode locations are defined by two uppercase letters followed by a number. The numbering system of electrode locations follows EEG convention where the locations on the left are odd-numbered and the locations on the right are even-numbered. The nomenclature of two uppercase letters is adopted to differentiate SC10-X/U labels from EEG system labels that use single uppercase letter (optionally followed by a lowercase letter) to identify the electrode location. The label letters for the electrode positions (SC10-X/U) are defined based upon the virtual vertical lines (figure 3) dividing the electrode space, symmetrical on either sides of the spinal cord. The electrodes are sequentially numbered for each line. Sequence of odd numbers for lines that reside on the left side, sequence of even numbers for lines that reside on the right side, and sequence of consecutive numbers for lines that reside in the centre.

The line that traces the anatomical midline of the spinal cord is defined as midline (ML) and encompasses the reference spinous processes; the label ML is used for the electrode locations on the midline. The virtual line named posterior line (PL) is defined by the electrode locations that divides the transverse curves in half on either side, label PL is used for the electrode locations on the posterior line. The virtual line that is defined along the left and right boundary of the inner-perimeter contour is referred to as lateral line (LL); hence, the label LL is used for the electrode locations on the lateral line. Label PM is used for the locations that reside between the posterior line and midline and label LP is used for the locations that reside between the lateral line and posterior line. These labels are defined by the combination of the letters from adjacent lines, where the first letter signifies the adjacent lateral line, and the second letter is defined by the adjacent medial line. Finally, the label AL is used to define the electrode locations that rest on AL and are in the region of C6 anterior tubercles.

### 2.4. Implementation of SC10-X/U using an Elastic Electrode Patch

An in-house non-invasive data recording elastic electrode patch was developed to realise the SC10-X/U electrode placement system. The electrode patch is designed to record spinal electrophysiological signals over spinal space covering cervical (neck) and upper thoracic (upper back) region up to T4 level. The designed elastic electrode patch can hold up to 76-channel electrodes for recording high-density electrospinography (ESG).

The electrode patch was realized in six sizes to incorporate the anthropometric (inter-individual) spinal cord length variability. The average inter-cervical spinous distance for each electrode patch is determined based upon the study by Ernst, M. J., et al. (Ernst et al., 2019); average inter thoracic spinous distance is based upon an earlier study (Ernst et al., 2013). The electrode patch was developed to offer: flexibility to accommodate for curvature around neck, shoulder, and back; vertical, and horizontal stretchability to accommodate inter-individual height and width variability; and comfort for long recording sessions.

The developed electrode patch is shown in Figure 6a. The extended lateral fins are used to hold the patch firmly against the neck (corresponding to the C4-C7 cervical spinal cord) using Velcro. The top part of the patch is held firmly against the participant with fastenings around the head. Elastic suspenders are used to hold the electrode patch around the ear and against the upper back (corresponding to the thoracic spinal cord). The electrode patch placed over the spinal cord of a participant can be seen in Figure 6b.

**Figure 6:**
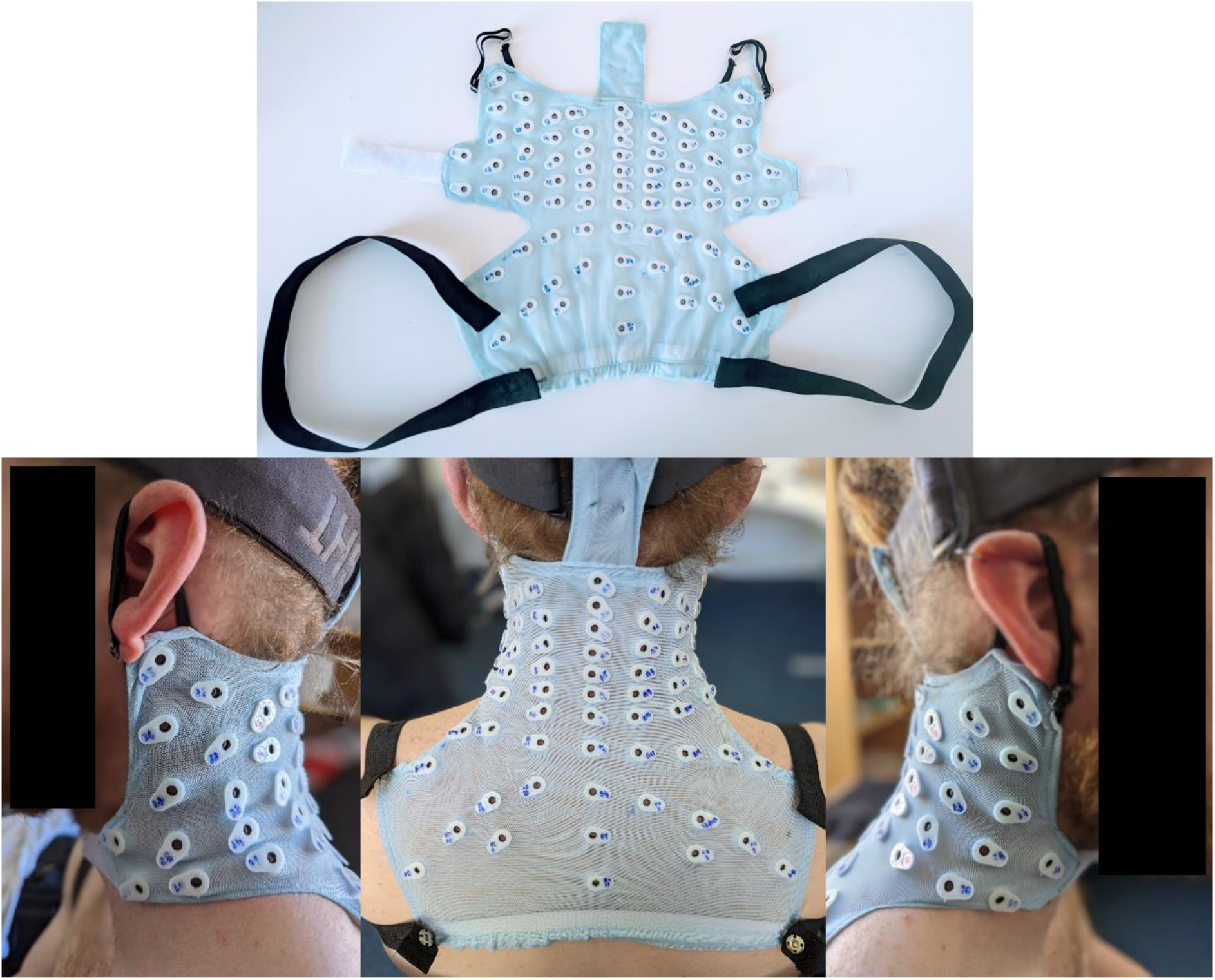
a. In-house developed electrode patch based upon the SC10-X/U electrode system. b. The electrode patch (up to 76 electrode channels) placed over the surface of the cervical and upper thoracic spinal cord space of an individual. *Note: The individual in the figure is a co-author and has consented to publicly share this figure*.

## 3. Experimental Demonstration of HD-ESG: Methodology

### 3.1. Participant and Ethics

Healthy participants (n=10, age: 28 ± 4, 4 Males) were recruited. The experiment protocol was explained to the participants and an informed consent (signed) was taken before the recording. Ethical approval was obtained from Tallaght Hospital/St. James’s Hospital Joint Research Ethics Committee for St. James’s Hospital, Dublin, Ireland (Project Reference: 0493).

### 3.2. Participant Exclusion Criteria

The exclusion criteria for this study include:

(i) History of neuromuscular, neurological, or active psychiatric disease.
(ii) Diagnosis of chronic cervical back pain or any other neck or upper back muscle stiffness, wear, and tear.
(iv) Recent history of severe backpain, suffering with spondylosis or spondylitis or any other severe back pain / spinal degeneration.

### 3.3. Experimental Design

The experimental part of the study was aimed at observing spinal potentials recorded using HD-ESG in response to peripheral stimulation via median nerve stimulation. In addition to HD-ESG, multichannel Electrocardiogram (ECG) signals were also acquired for removing ECG artifact from the recorded ESG signals.

During the experiment, the participants were instructed to lean forward and sit comfortably in an inclined position on the physio-chair as shown in Figure *7*b. In this position, the legs formed approximately a 60-degree angle at the knee, and the chest rest was positioned at roughly 30 degrees from the vertical plane. Throughout the recording session, the participant’s head rested on the chair’s headrest, and their hand was placed on the handrest, forming a 90-degree angle at the elbow. Participants were instructed to keep their eyes closed during the experiment.

**Figure 7:**
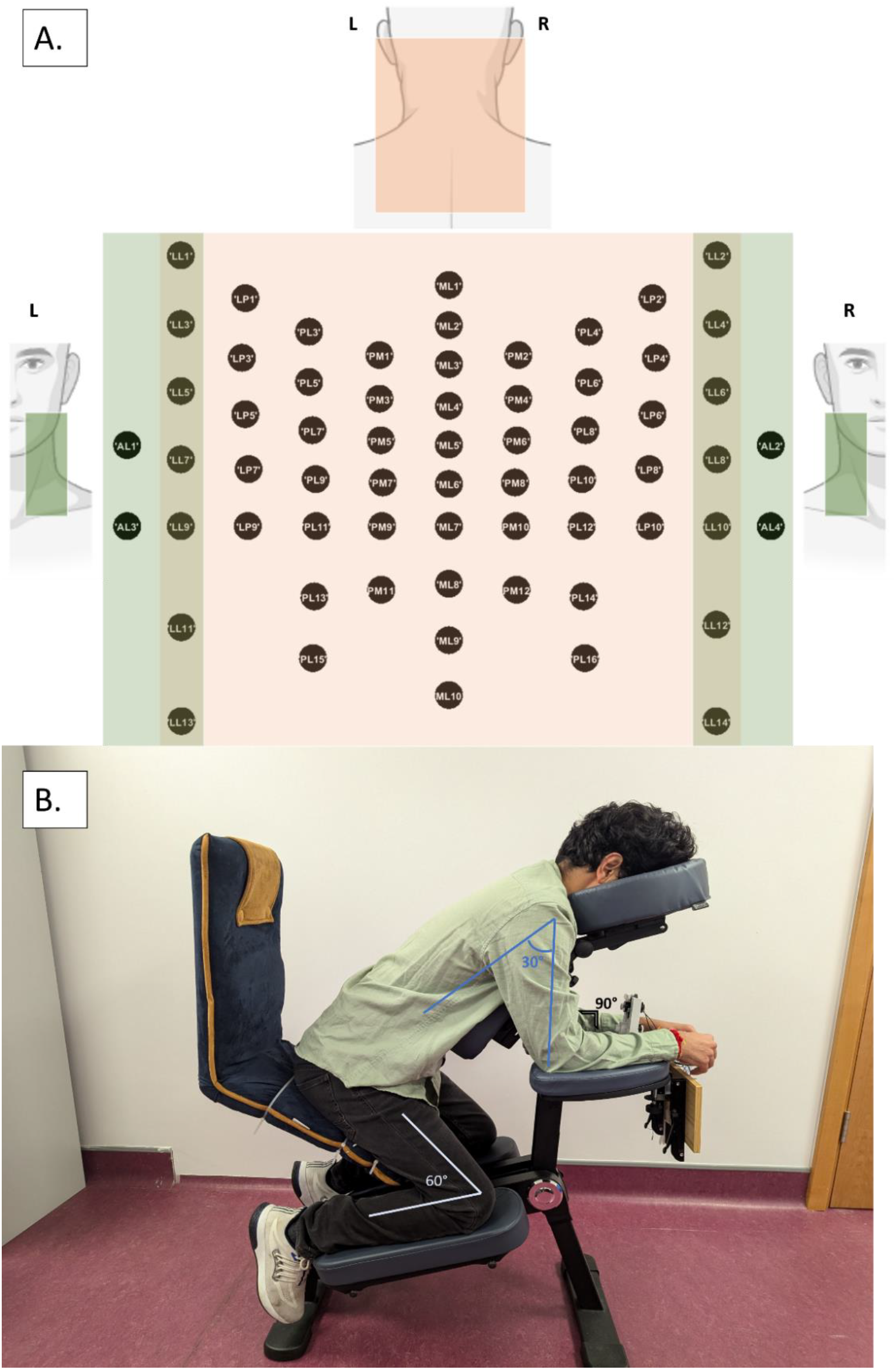
A. The 64-electrode channels used for the recording and its corresponding electrode space. B. A demonstrator sitting in an inclined position on the physio-chair. The demonstrator’s chest was supported by the chest rest while their head rested on the headrest and hands rested on the handrest respectively. *Note: The demonstrator in the figure is a co-author and has consented to publicly share this figure*.

The experiment consisted of a single session of median nerve stimulation; the session lasted for approximately 20 mins. Each session consisted of 35 blocks, each block lasted for 30 seconds and consisted of 20 seconds of stimulation followed by 10 seconds of rest. In the stimulation phase, a train of stimulation pulses at an average frequency of 2 Hz were applied at the wrist of the right hand to elicit stimulation of the median nerve. Stimulation was applied using a constant current stimulator DS7A (Digitimer Ltd., UK). DS7A is MDD CE medically certified and can provide up to 100mA constant current high voltage pulses. The stimulation intensity was set to 1.5 times the motor threshold and the pulse width was set to 200µs. The motor threshold was defined as the minimum intensity at which prominent thumb movement was observed with consistency. In the session, a total of 1400 trial responses (35 block x 40 stimulations per block) to peripheral stimulation were recorded.

#### Data acquisition

The data collection system consisted of a high-density biosignal amplifier that was used for high density ESG, and multi-channel ECG. The electrical activity of 64 (ESG) + 5 (ECG) + 8 (EXG) channels were recorded simultaneously using the high-speed Biosemi-ActiveTwo system (Biosemi B.V., Amsterdam, The Netherlands) at a sampling frequency of 8 kHz, with a bandpass filter over 0Hz - 1600Hz frequency range. All electrode channels were visually inspected for signal quality prior to recording and the electrode offset was kept between ± 25mV.

The HD-ESG signals were recorded using 64-channel active electrodes with placement following the SC10-X/U electrode system. The 64-channel ESG montage was derived from the proposed system (Figure *7*a). The active electrodes were connected to the electrode holder and electrolyte gel was used to enhance skin-electrode connection.

Multi-channel ECG was recorded in accordance with 5-electrode ECG system with flat active sintered Ag-AgCl electrodes. The following 7-lead ECG signals were extracted from the ECG system:

1. *Lead I* = *LA* − *RA*
2. *Lead II* = *LL* − *RA*
3. *Lead III* = *LL* − *LA*
4. 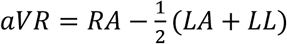
5. 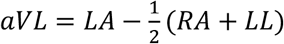
6. 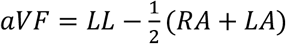
7. 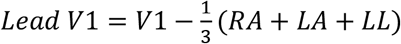

Eight additional signals were collected using flat active sintered Ag-AgCl electrodes. The electrodes were placed at the following locations: (i) left and right earlobe, (ii) to the left of the left eye and to the right of the right eye, (iii) left and right clavicle, (iv) below left eye, (v) er ‘s p int (brachial plexus).

### 3.4. Data Analysis

The signals were analysed using MATLAB (The MathWorks, 2023) scripts in conjunction with the FieldTrip toolbox (Oostenveld et al., 2011) and the NoiseTool toolbox (de Cheveigné et al., 2018).

#### 3.4.i. Signal pre-processing

Two main contaminating artifacts were identified in the recorded ESG signals: stimulation and ECG artifacts. The stimulation artifact lasted approximately 1-2ms; this artifact was identified and the signal during this period was interpolated using piecewise cubic interpolation. ECG artifact from the ESG signal was removed by applying canonical correlation analysis (CCA) (de Cheveigné et al., 2018; Hotelling, 1936). This was achieved by isolating and removing the two canonical components that represented the highest correlation between ESG data (64 channels) and ECG leads (7 leads). The first two canonical components (CC) were selected for removal, as the correlation (r) score after first two CC’s demonstrated sharp decrease in magnitude.

The remaining CC’s were back-transformed to the original space to provide the denoised ESG signal. The signal post artifact rejection was re-referenced using ‘common average referencing’. The re-referenced signals were baseline corrected and segmented into epochs of 0.5 seconds, starting 0.1s before the stimulation until 0.4s after the stimulation. Before further processing of the signal, bad channels were visually determined and interpolated (weighted average) based on the signal from neighbouring channels (neighbour distance <=50mm). The epoched trials were visually inspected and the bad trials were rejected. The remaining trials (1287 ± 91 trials, ∼92%) were bandpass filtered between 10 and 1500 Hz (Akaza et al., 2020), dual-pass Butterworth filter.

#### 3.4.ii. Event-Locked Activity

The pre-processed epochs from the ESG channels underwent event-locked analysis. The epochs were averaged over all the good trials. Surface Laplacian referencing, based on Hjorth’s method (Carvalhaes and de Barros, 2015; Hjorth, 1975) was applied to the averaged signals. The resulting signals were analyzed to identify characteristics of the event-locked (time-phase locked) evoked activity and spatial distribution of the evoked activity.

### 3.5. Statistical Analysis

A statistically significant difference from zero of the evoked activity was identified at each time of the recorded signal from all participants across all trials, using the Wilcoxon signed-rank non-parametric test (Iyer et al., 2017; Nasseroleslami et al., 2014). To account for multiple comparisons in 64 channels and several time points (-0.001s - 0.03s), we retained the dominant 4 principal components that explained 85% of the total variance, plus an additional component (4+1=5). Bonferroni correction was subsequently applied at α = 0.01.

## 4. Demonstration of HD-Electrospinography: Results

High-density electrospinography (HD-ESG) using non-invasive 64-channel surface electrodes was used to record spinal activity in response to median nerve stimulation. The event-locked activity was evaluated for the recorded signals.

### 4.1. Average Evoked Potential

The average (across trials) evoked potential for individual participants and the grand-mean signal across participants at channel ML6 (C6 vertebral level) are shown in Figure 8 (respectively in grey and black). Significant evoked potentials P9 and N13 were recorded at ML6 (C6 cervical level); marked in red in Figure 8. The observed latency of the evoked potential N13 and P9 across individual participants was 13.2 ± 1.1ms (mean ± sd) and 9.7 ± 1.2ms (mean ± sd) respectively.

**Figure 8:**
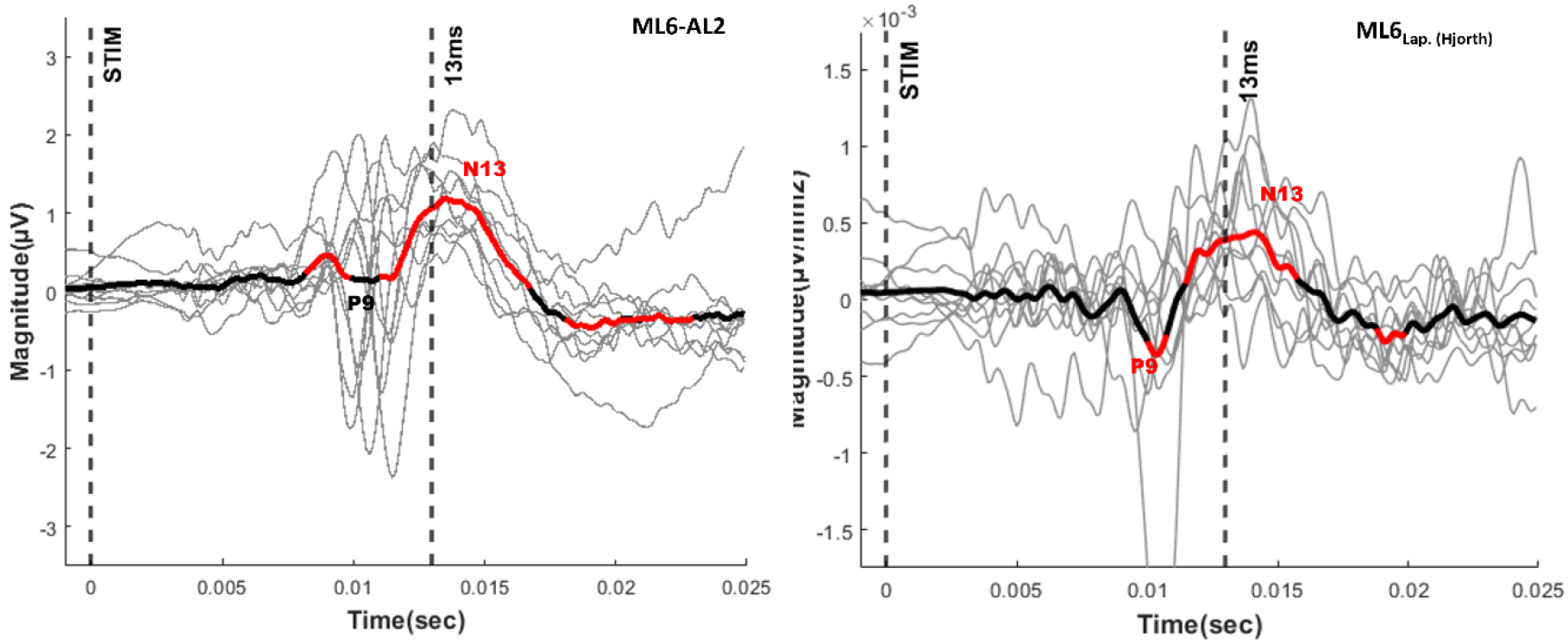
Grand-mean of the evoked potentials across all participants at ML6 electrode. Grey lines represent individual participant evoked potentials, and thick black line correspond to the grand mean evoked potential across all participants. The duration of significant evoked response (p<0.01)is highlighted by red lines. The figure indicates significant N13 and P9 evoked potentials. 6a. ML6 response in reference to AL2 electrode (left). 6b. surface Laplacian re-referenced ML6 response (right).

Electrode channels AL1, LL7, LP7, PL9, PM7, ML6, PM8, PL8, LP8, LL8, and AL2 from the SC10-X/U system are comparable to the traditional ring electrode placement at the C6 vertebral level traditionally used to record spinal evoked potential in response to median nerve stimulation (Restuccia and Mauguiére, 1991). The grand mean signal across participants for the channels used to define the ring electrodes system is shown in Figure 9. In Figure 9, the evoked potential at 13 ms has the highest amplitude at channel ML6 and reverses polarity at the anterior-lateral electrode channel (AL2).

**Figure 9:**
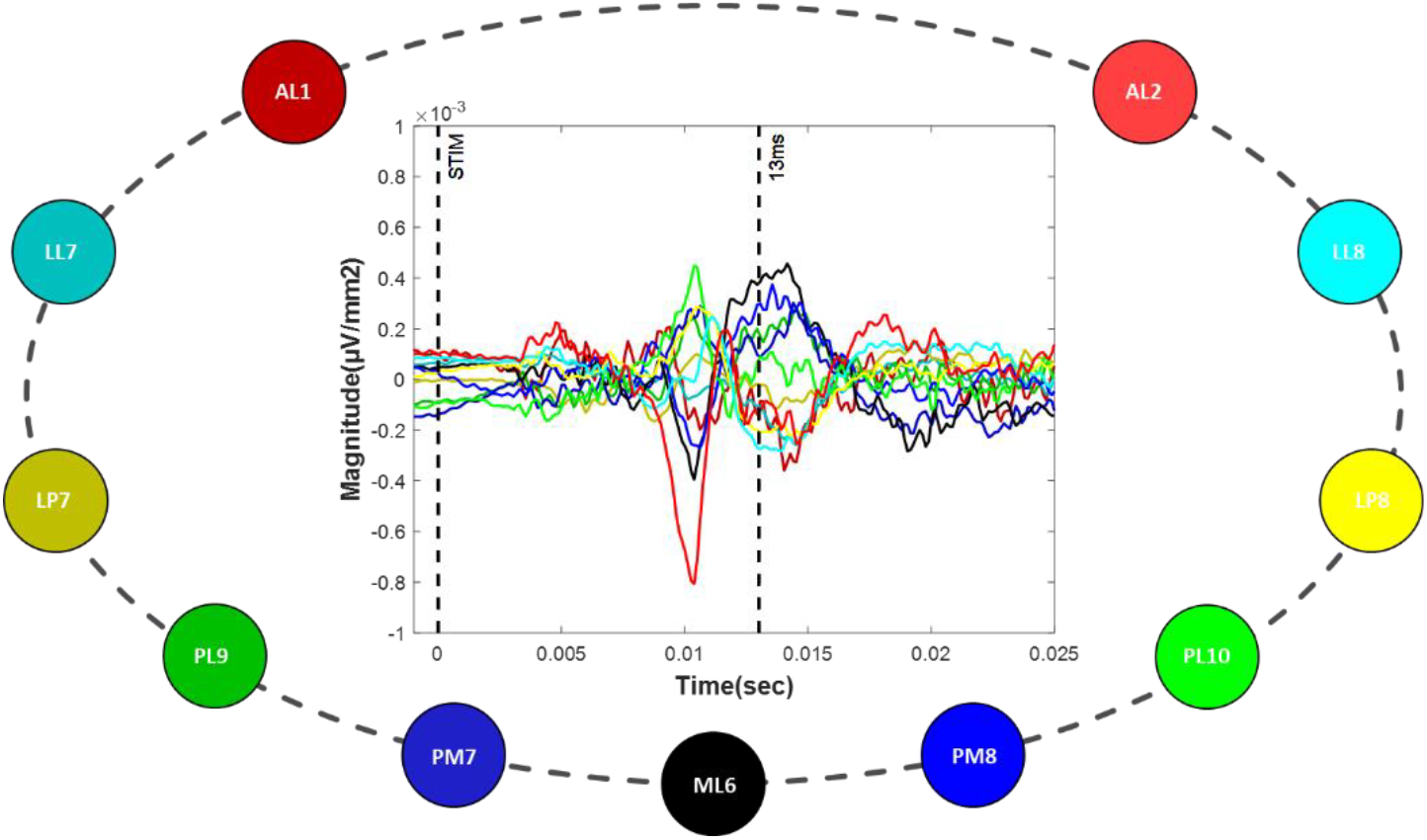
Grand-mean signals plotted for SC10-X/U channels at Cv6 compatible with ring electrode placement system at Cv6. (Surface Laplacian re-referenced)

### 4.2. Topographic Map of Evoked Spinal Potentials

High density spinal activity recorded using the SC10X/U electrode placement system was used to map the spatial distribution of the electrical activity across the spinal space. The topographical plot of grand-mean spinal evoked response during the time frame 12-14ms (N13) after the median nerve stimulation is shown in Figure *10*. The region around the midline indicates higher spinal response activity with its epicentre at channels ML4-ML6. Another epicentre of spinal response with reversed polarity is also indicated in Figure *10* at the lower cervical right-anterior-lateral and right-lateral regions.

The grand-mean response of the midline electrodes in the cervical region (ML2-ML7) can also be seen in Figure 10. The lower cervical level channels on the right side of the Topography map (Figure 10) indicate reversed polarity of the evoked potentials in the region.

**Figure 10:**
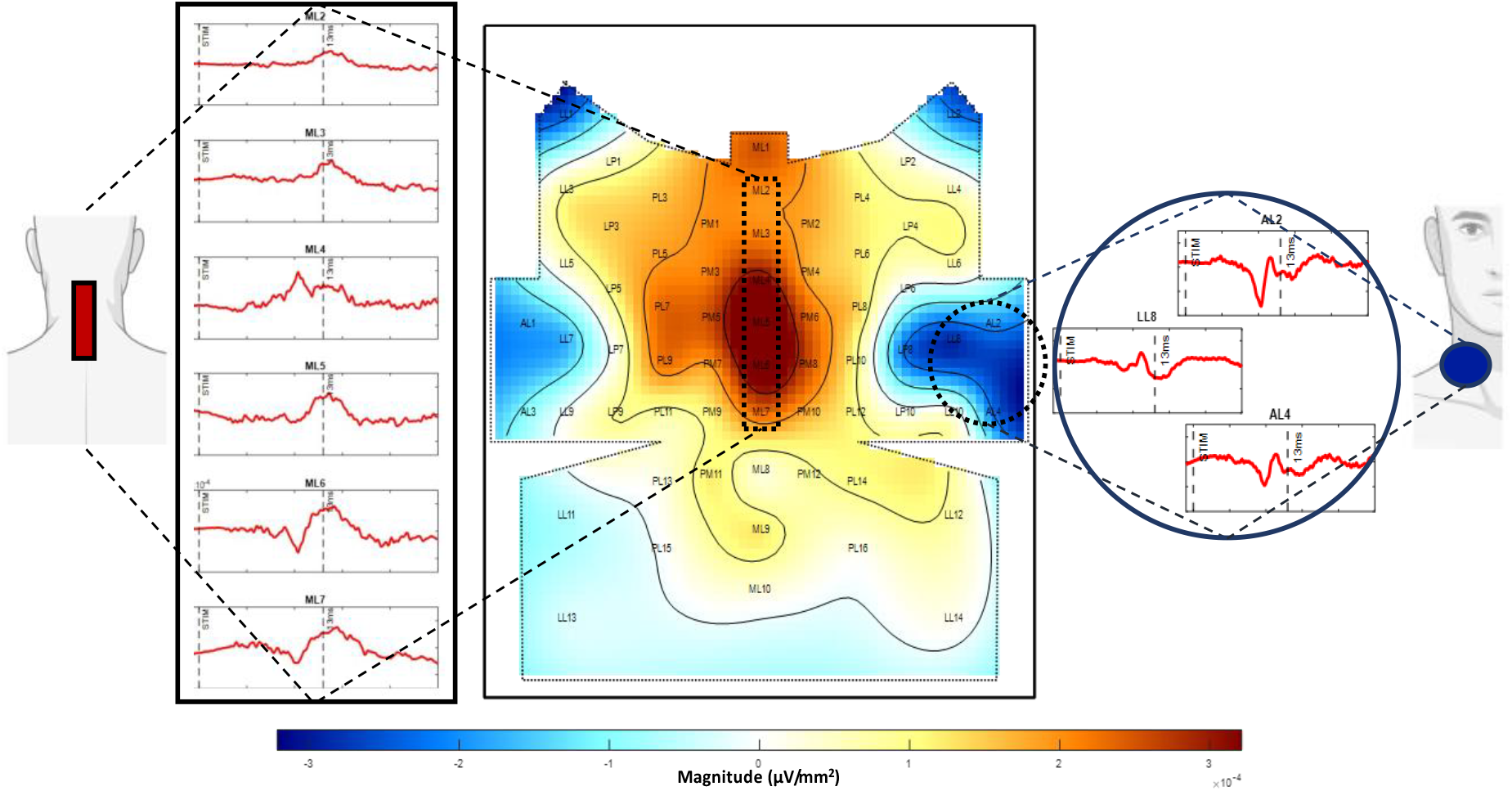
Topographic Map of Evoked Spinal Potentials. Centre Figure. Topographic Map of the grand-mean spinal evoked response over the SC10-X/U sensor space, time-window (12ms-14ms). Left figure: corresponds to the spinal activity recorded from channels ML2-ML7 (midline, posterior) for the time frame between 0.001s before stimulation and 0.025s after stimulation. Right figure: corresponds to the spinal activity recorded from lateral-anterior channels for the time frame between 0.001s before stimulation and 0.025s after stimulation.

## 5. Discussion

This study proposes an HD electrode placement system (SC10-X/U) for recording the electrophysiology activity of the cervical spinal cord. The utility of the proposed SC10X system was demonstrated by recording evoked spinal activity (HD-ESG) in response to median nerve stimulation with high spatial resolution. The SC10X/U system promotes a standardized methodology for recording high-density ESG signals at the cervical and upper thoracic spinal levels. This addresses the technical gap leading to non-standardized multichannel recordings being utilized by recent studies for estimating spinal signals in response to external stimulation (Akaza et al., 2020; Mardell et al., 2022; Sumiya et al., 2017) or to develop spatial filters for accessing single trial spinal evoked potentials (Nierula et al., 2022). These studies indicate the potential benefits of high-density spinographic techniques over non-HD recordings but fail to facilitate compatible and reproducible research across individuals and research groups. The proposed system maps the cervical and upper thoracic region to the electrode space for non-invasive neuroelectric imaging and can incorporate up to 76 channels as shown in Figure 3. This is especially suitable for sensorimotor studies investigating the upper extremities.

Development of the SC10-X/U electrode placement system is aimed at the standardization of future studies, to enable compatible, comparable, and reproducible methods across individuals and research centres. EEG electrode placement systems (10/20, 10/10, 10/5) have proven to be an invaluable tool in the field of neuro-electro-physiology, enabling standardized electrophysiological recording resulting in applications including development of biomarkers (de Aguiar Neto and Rosa, 2019; McMackin et al., 2019), brain-computer interface (Lotte et al., 2007; Zhang et al., 2018), and resting-state analyses (Dukic et al., 2022; Metzger et al., 2023). Similarly, standardized high-density electrospinography presents huge potential in uncovering the role of the spinal cord in sensorimotor processing within the central nervous system and moving away from treating SC as a “wire” relay between the brain and periphery.

The SC10-X/U system was demonstrated using a 64-channel electrode patch to record evoked spinal activity in response to the median nerve stimulation. Short latency SEP’s P9, and N13 were recorded as shown in Figure 8, and corroborate previous studies (Cruccu et al., 2008; Mauguiére, 2003) with consistent SEP measures and anatomically valid location of responses that represented the corresponding dermatome (Cramer et al., 2014). Posterior cervical channels showed a negative evoked response approximately 13ms (N13) after stimulation. The N13 potential is believed to be generated by the dorsal horn and to have a post-synaptic origin (Desmedt and Cheron, 1981; Jeanmonod et al., 1989; Urasaki et al., 1990). We also observed reversed polarity of N13 response when moving from the posterior (ML6) to right-anterior electrodes (AL2) at an approximate C6 vertebral level, as demonstrated in previous studies (Desmedt and Huy, 1984; Restuccia and Mauguiére, 1991).

The spatial distribution of the observed spinal activity was mapped using the HD electrode channels. The topographic map indicated that high cervical spinal activity is localized around the midline with its epicenter at ML4-ML6 level (Figure 10). This epicentre could correspond to fibres from roots of spinal nerves C5-C7, innervation of which is contained in the median nerve. Moreover, an epicenter of high activity was also seen around the anterior and lateral region at lower cervical levels, through the polarity is reversed. The observed evoked potentials and their characteristics are compatible with the existing literature (Cruccu et al., 2008; Mauguiére, 2003). The results from previous studies recorded using ring electrode placement (Restuccia and Mauguiére, 1991) can be extended and interpreted using the comparable electrode locations from the proposed SC10-X/U system, i.e. electrode channels AL1, LL7, LP7, PL9, PM7, ML6, PM8, PL8, LP8, LL8, and AL2 from SC10-X/U system are comparable to the traditional ring electrode system. Here, we demonstrated in Figure 9 that the evoked potentials recorded at the selected electrode channels are comparable to the evoked activity reported in studies using ring placement electrode system (Bankim Subhash Chander et al., 2022; Restuccia and Mauguiére, 1991).

Development of the standardised HD-ESG recording system will enable analysis of evoked responses using combined spatial, spectral, and temporal techniques similar to those used for HD-EEG. Source localization techniques (Akaza et al., 2020; Sumiya et al., 2017) will also improve the understanding of spinal activity and allow further investigation of dysfunction of spinal cord in neurological conditions.

The future direction of this study would be to investigate the spatiotemporal changes in the evoked HD-spinal potentials in response to peripheral nerve stimulation in neurodegenerative conditions, such as amyotrophic lateral sclerosis, that affect sensorimotor communication.

## 6. Conclusion

The SC10-X/U electrode system described in this paper provides full coverage of the cervical and upper thoracic region by closely and evenly spaced high-density electrodes applied at standardized locations with respect to accessible anatomical locations. The system can incorporate up to 76 electrode positions, though certain studies may require only a subset of electrodes to cover the spinal space of interest. The utilization of this system in studies of spontaneous (such as resting state, tast-related) and evoked ESG activity is expected to facilitate compatible and reproducible research. In theory, information derived from the surface-recorded ESG could be projected onto the anatomical space and communicated through the use of a standardized atlas across different research and participant groups, promoting the identification of sources of neural activity within the spinal cord. This should promote standard practices across large-scale neurophysiological and clinical studies of neurological conditions.

